# Functionality of potato virus Y coat protein in cell-to-cell movement dynamics is defined by its N terminal region

**DOI:** 10.1101/2025.04.29.651159

**Authors:** Anže Vozelj, Tjaša Mahkovec Povalej, Katja Stare, Magda Tušek Žnidarič, Katarina Bačnik, Valentina Levak, Ion Gutiérrez-Aguirre, Marjetka Podobnik, Kristina Gruden, Anna Coll, Tjaša Lukan

**Author notes:** shared last authorship (Anna Coll, Tjaša Lukan).

## Abstract

Potato virus Y (PVY) is one of the top ten economically most important plant viruses and responsible for major yield losses. We previously suggested the involvement of the N terminal region of potato virus Y coat protein (CP) in PVY spread. By constructing different N terminal deletion mutants of PVY N605 strain, we here show that deletions of 40 or more amino acid residues from the N terminal region of the CP resulted in the PVY multiplication limited to primary infected cells in *Nicotiana clevelandii* plants. Deletion of 26 residues profoundly impaired PVY cell-to-cell movement and prevented systemic PVY spread, while deletions of 19-23 residues allowed delayed systemic PVY spread. Introduced point mutations in the identified region prevent (S21G) or delay (G20P) PVY movement. In summary, this work shows the significance of the CP N-terminus for movement of the PVY.

## Introduction

Plants undergo constant exposure to various pathogens and pests, with viruses being the most severe amongst them. Potato virus Y (PVY) is classified amongst the top ten economically most important plant viruses infecting solanaceous crops including potato, tomato, tobacco and pepper (1). PVY is the causal agent of potato tuber necrosis ringspot disease (2), a devastating disease that negatively impacts potato tuber quality and yield (3).

PVY belongs to the *Potyvirus* genus. Its genome consists of 9,7-kb positive sense single stranded RNA (+ ssRNA) encoding for single open reading frame (ORF) that is translated into 350 kDa long polyprotein which is cleaved into 10 mature proteins (3). Furthermore, PVY genome contains an additional ORF *pipo* (“pretty interesting *potyviridae* ORF”), embedded within the P3 cistron of the polyprotein. Expression of *pipo* occurs through transcriptional slippage by the viral RNA polymerase and results in production of the P3N-PIPO fusion protein, which is involved in cell to cell movement (4, 5).

The majority of potyviral proteins are multifunctional and they altogether contribute to the establishment of successful viral infection, that encompasses multiple steps including transmission by aphids, penetration of viral particles into the cell, virion disassembly, translation, replication, suppression of host defense mechanisms, virion assembly and virus movement from the primary infected cells to neighboring cells and systemic spread (6).

Viral movement is facilitated through plasmodesmata, plant specific structures connecting two adjacent cells that serve as gateways for cell-to-cell movement and contribute to the establishment of systemic infection (7, 8, 9, 10). At least four potyviral proteins, coat protein (CP), cylindrical inclusion protein (CI), P3N-PIPO and the second 6 kDa protein (6K2) protein are essential for cell-to-cell movement of potyviruses (11). P3N-PIPO is localized in the plasmodesmata and directs CI to form conical structures that are crucial for assistance of potyviral intercellular movement through plasmodesmata, which was shown for turnip mosaic virus (TuMV) in *Nicotiana benthamiana* (12). Accumulation of CP in plasmodesmata during infection is also crucial since together with the potyviral Helper-Component Proteinase (HCPro) increases the size exclusion limit (SEL) of plasmodesmata, which was determined by microinjection studies for bean common mosaic necrosis potyvirus (BCMNV) in *N. benthamiana* (13). Systemic infection encompasses viral movement through phloem and less often through xylem to reach and infect plant distant tissues (14, 15). However, before reaching vasculature, plant viruses need to cross various cellular barriers including bundle sheath, vascular parenchyma and companion cells, to reach sieve elements, through which they are transported to distant tissues (14). The invasion of distant tissues requires unloading from sieve elements into companion cells and cell-to-cell movement into bundle sheath and mesophyll cells (14). Despite PVY being one of the most extensively studied potyviruses on molecular scale, information about crucial factors governing cell-to-cell as well as systemic spread are scarce (6, 11).

The cryo-EM structures of some potyviruses including PVY, watermelon mosaic virus (WMV) and TuMV virions have been determined and offered a detailed insight into the conserved helical arrangement of CPs assembled around viral ssRNA as well as the conserved three dimensional structure of CP among them (17, 16, 18). CP is the only potyviral structural protein and more than 2000 copies of CP encapsidate the viral ssRNA molecule to form potyviral virions (6, 10). Individual CP is built of three distinct regions. The N terminal region with residues up to Val44 in PVY, which are exposed at the outer surface of the virus, are flexible and thus its 3D structure was not determined. The rest of N terminal region, from Val44 to Gln76, is structured. It has an extended structure with one short alpha helix in the middle (17). The central core has a globular shape. The C terminal region is in the lumen of the virus and its extended structure is supported by the viral ssRNA scaffold (17). Beyond its primary role in viral encapsidation, the CP is indispensable for aphid transmission, potyviral RNA amplification and cell-to-cell movement (10, 13, 19). Functional studies of the CP protein indicate crucial roles of its N and C terminal regions and the core region in potyvirus infectivity. In the case of PVY, the absence of the flexible CP N terminal region hinders the formation of filaments and cell-to-cell viral movement (17). For soybean mosaic virus (SMV), the CP C terminal region is involved in CP intersubunit interactions, viral cell-to-cell and long distance movement and virion assembly (20). In potato virus A (PVA), phosphorylation of Thr243 residue on CP C terminal region is crucial for PVA replication *in planta* (21). These results and results of several other studies suggest that different CP regions are important for cell-to-cell movement of potyviruses (22, 23, 24, 25, 26, 27). It is not known yet in what structural shape are the viruses propagated through the plant, as assembled virions or viral ribonucleoprotein (RNP) complexes associated with CP (14, 15).

The so far generated evidence-based knowledge differs from one potyvirus to another, and thus, more research is needed to comprehensively elucidate these complex processes, particularly for those economically relevant potyviruses such as PVY. By constructing different PVY N terminal deletion mutants, we identified the regions of the CP that are important for efficient PVY cell-to-cell movement.

## Results

### Deletion of 40 or more amino acid residues from the PVY CP N terminal region affects cell-to-cell movement, but does not prevent replication

Previously we showed that PVY with deleted 50 N terminal amino acid residues was still able to replicate but based on the low and spatially limited levels of RNA accumulation, we suggested that N terminal region is required for cell-to-cell movement (17). To confirm this hypothesis, we here constructed two GFP-tagged PVY (PVY N605-GFP) mutants lacking either 40 or 50 amino acid residues at the CP N terminal region (hereafter ΔN50-CP and ΔN40-CP). In all constructed mutants GFP was inserted between NIb and CP coding sequence, flanked by protease recognition sites, allowing GFP excision from the polyprotein after translation (28). We followed the spread of ΔN50-CP and ΔN40-CP with confocal microscopy upon bombardment of *Nicotiana clevelandii* leaves (Fig. 1B). Neither mutant was able to move cell-to-cell, remaining limited to bombarded cells (Fig. 1B). We detected viral RNA replication of both mutants in the bombarded leaves, albeit at significantly lower levels in comparison to the non-mutated clone (hereafter WT-CP) (Fig. S1, data S1 and S2). Despite the lower RNA levels observed in leaves bombarded with constructed mutants, confocal microscopy image analysis showed that the accumulation of GFP in individual infected cells was the same in WT and ΔN40-CP mutant, confirming unhindered replication (Table S2). These findings corroborate our previous results showing that N terminal region CP truncations still allow viral replication (17) and further show that the absence of either 50 or 40 N-terminal residues abolish cell-to-cell viral movement.

**Figure 1.**
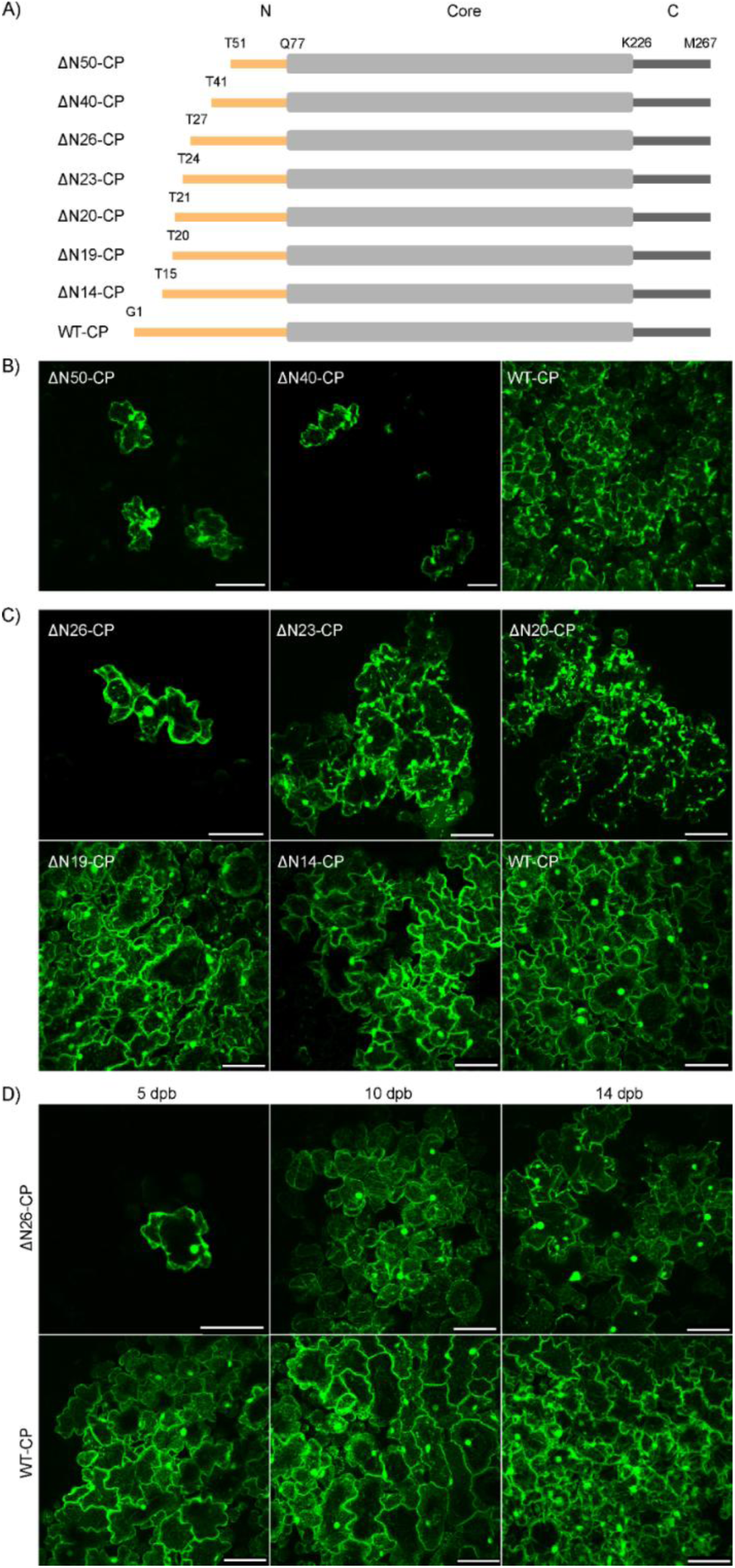
Amino acid deletions at PVY CP N terminal region influence viral cell-to-cell movement. **(A)** Schematic representation of constructed mutants with deletions on PVY CP N terminal region (N-terminal region, orange; core region, grey; C-terminal region, dark grey) with labeled amino acids. **(B)** Confocal microscopy images showing viral replication is limited to single cells when 50 (ΔN50-CP) or 40 (ΔN40-CP) amino acid residues were deleted at CP N terminal region, while non-mutated clone (WT-CP) shows cell-to-cell spread 10-12 days post bombardment (dpb). Scale bar: 100 µm. **(C)** 5 dpb viral replication is limited to single cells in ΔN26-CP mutant, while cell-to-cell virus spread was observed for CP deletion mutants ΔN23-CP, ΔN20-CP, ΔN19-CP, ΔN14-CP. Scale bar: 100 µm. **(D)** ΔN26-CP cell-to cell-spread was observed at later time points (10 and 14 dpb). Scale bar: 100 µm.

### The deletions of 19-26 amino acid residues of the PVY CP N terminal region are influencing dynamics of viral cell-to-cell spread

To further assess which region of CP N terminal is crucial for cell-to-cell virus spread, we constructed GFP-tagged mutants lacking 26, 23, 20, 19 and 14 amino acid residues at the CP N terminal region (Fig. 1A) and followed their localization under confocal microscope at different time points after virus inoculation (Fig. 1, C and D, data S2 and Table S1). Mutants lacking 26 (ΔN26 - CP) and 23 (ΔN23-CP) residues on CP N terminal region had the first non deleted amino acid replaced with glycine (G) during the mutagenesis.

At 5 dpb, ΔN14-CP and ΔN19-CP mutants did not show any differences in cell-to-cell spread compared to WT-CP, while a lower spread level was noticed for ΔN20-CP and ΔN23-CP (Fig. 1C). The most pronounced difference in comparison to other mutants and to WT-CP was observed in the case of ΔN26-CP mutant, which remained confined to single cells at 5 dpb (Fig. 1, C and D). However, at later timepoints (10 and 14 dpb), in approximately half of the infected plants, we also observed cell-to-cell viral movement, although substantially delayed compared to WT-CP (Fig. 1D and Table S1). Also, the measured viral RNA load for ΔN26-CP mutant was significantly lower than that of WT-CP at all 5 dpb, 10 and 14 dpb (Fig. S2 and data S1).

Whole plant imaging was conducted to more precisely examine the dynamics of cell-to-cell virus spread of WT-CP, ΔN23-CP, ΔN19-CP and ΔN14-CP through time on inoculated *N. clevelandii* leaves (Fig. 2A). As expected, the results revealed statistically significant differences in the viral multiplication area between N23-CP and WT-CP, and ΔN19-CP compared to WT-CP (Fig. 2, B and C and data S3 and S4). A similar though less evident trend was observed in the case of ΔN14-CP compared to WT-CP, which was not statistically significant (Fig.2 C and data S4).

**Figure 2.**
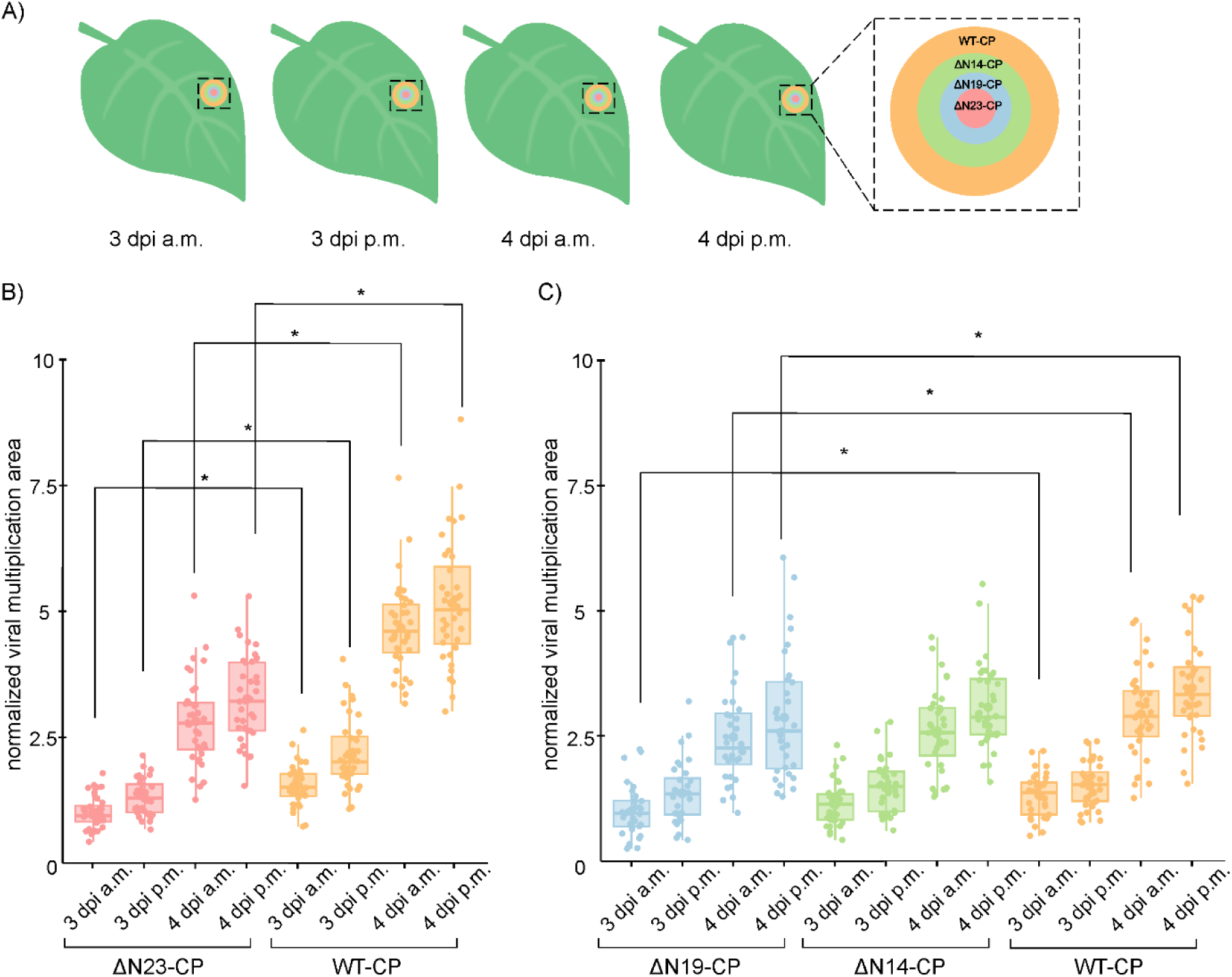
Precise analysis of the viral multiplication areas revealed a hampered cell-to-cell virus spread in ΔN19-CP and ΔN23-CP mutants. **(A)** The scheme of the viral multiplication area at different time points for all mutants compared with WT-CP. The area was followed by more precise analysis using whole plant imaging system for examining cell-to-cell virus spread of ΔN23-CP, ΔN19-CP, ΔN14-CP and WT-CP in four timepoints including 3 days post inoculation (dpi) in the morning (a.m), 3 dpi in the afternoon (p.m), 4 dpi a.m. and 4 dpi p.m. Note that the viral multiplication area was the largest after infection with WT-CP and decreased in the latter order ΔN14-CP, ΔN19-CP and ΔN23-CP. **(B and C)** Normalized viral multiplication area. After inoculating leaves with WT-CP or mutants, plants were imaged with whole plant imaging system. Images of three selected viral multiplication areas per leaf per plant were taken at four time points. Differences were statistically evaluated using Welch’s t test. Results are presented as boxplots with dots representing normalized viral multiplication areas (data S3 and S4) for ΔN23-CP-CP, WT-CP **(B)** and ΔN19-CP, ΔN14-CP, WT-CP **(C)**. Statistically significant differences (p < 0.05) are denoted with an asterisk (*). Vertical lines present all points except outliers. Raw and normalized data, number of plants and statistics are specified in Data S3 and S4. The same trend was observed in a replicate experiment (Fig. S3 and data S5). For comparison between two independent experiments, data measured for mutant viruses were normalized to median of values obtained for WT virus (data S6).

These results show that the dynamics of viral cell-to-cell spread is dependent on the length of the CP N terminal region (Fig. 1 and data S2). More precisely, as the number of CP N terminal region deleted amino acid residues increases, cell-to-cell movement speed is decreased (Fig. 2, B and C and data S3, S4 and S6). We conclude that amino acids in the region 19-26 on CP N terminal region are essential for an optimal PVY cell-to-cell movement. Furthermore, with electron microscopy we observed that viral assembly at all tested deletion mutants is feasible (Supp. Fig. 10).

### Systemic viral spread dynamics is affected by the cell-to-cell movement

Since viral cell-to-cell spread is prerequisite for systemic viral spread, we further tested if the observed delay in cell-to-cell viral spread of mutants (Figs. 1, C and 2, B and C) affects the systemic viral spread. In contrast to the systemic spread of WT-CP, which is detectable already at 7 dpb, the spread of ΔN23-CP, ΔN19-CP and ΔN14-CP mutants to systemic leaves occurs with a delay (Fig. 3A). If compared to WT-CP, ΔN14-CP was observed in systemic tissue with 1 day delay (8 dpb), ΔN19-CP with 2 days delay (9 dpb) and ΔN23-CP with approximately 3 weeks delay (25 dpb) (Fig. 3A). The mutant ΔN26-CP did not reach systemic leaves even at the highest tested dpb (Fig. S4).

**Figure 3.**
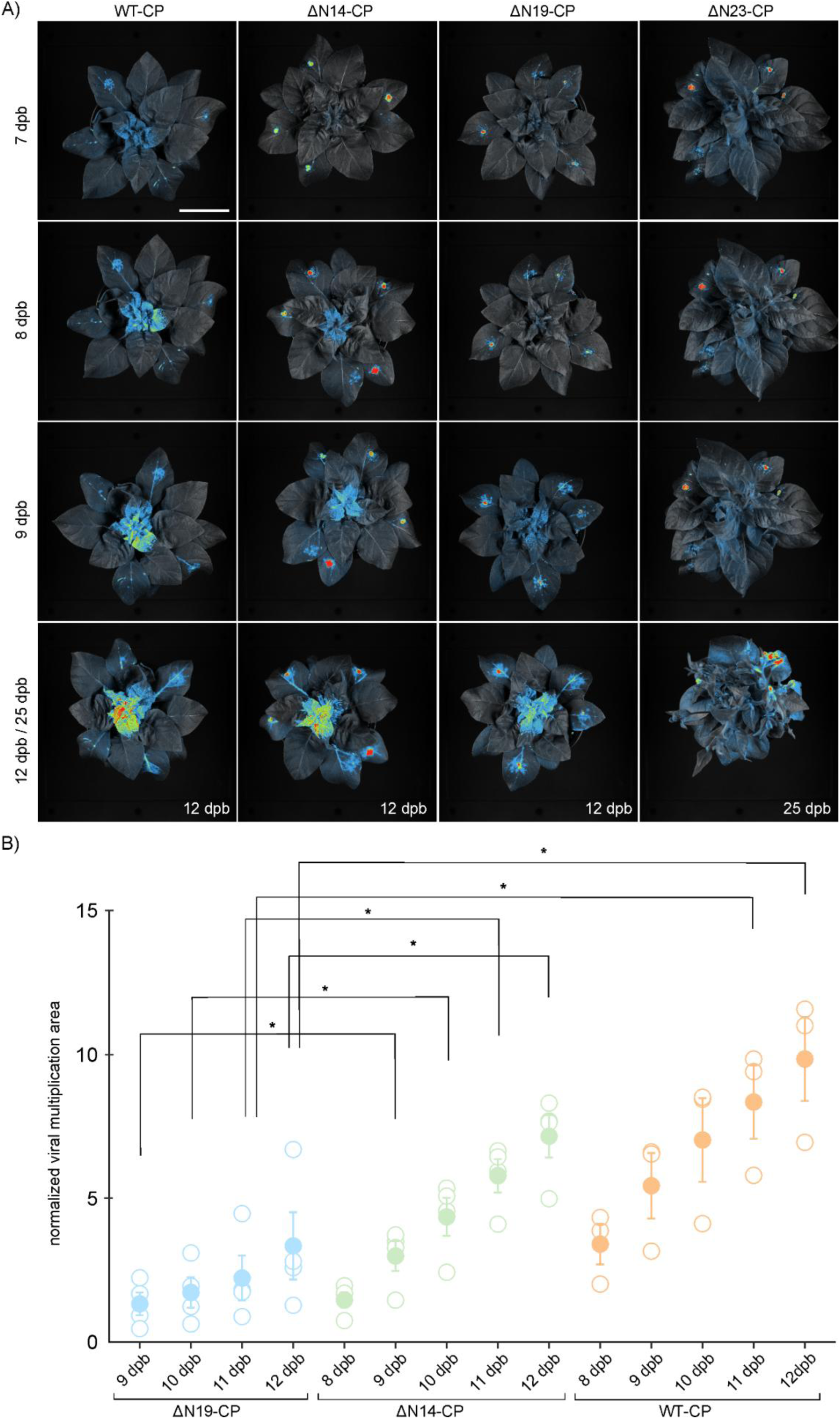
Systemic viral spread of mutants is delayed similarly as cell-to-cell movement. **(A)** Spatio-temporal PVY distribution in *N. clevelandii* systemic tissue. Viral multiplication area was followed 7-12 and 25 days post bombardment (dpb), by using whole plant imaging system. Scale bar: 5 cm. **(B)** Quantification of viral multiplication area in ROI in the *N. clevelandii* systemic tissue in total count per mutant in five selected timepoints including 8-12 dpb. Since ΔN19-CP did not spread systemically at 8 dpi, normalized viral multiplication area is not shown. Mean (represented with filled dot) and standard error are shown.

Individual measurements are shown as empty dots, representing normalized viral multiplication area. Differences were statistically evaluated using Welch’s t test. Statistically significant differences (p < 0.05) are marked with asterisk. Raw and normalized data, number of plants and statistics are specified in data S7. Since ΔN26-CP is not capable of spreading systemically, it was not included in the analysis. Also ΔN23-CP had such a delayed spread that it was not possible to do a quantification of viral multiplication area at 8 to 12 dpb (Fig. 3A).

Quantitative analysis of viral multiplication signal in systemic tissue confirmed that WT-CP achieved the largest multiplication area, followed by ΔN14-CP and lastly ΔN19-CP in all observed time points (Fig. 3 and data S7). When comparing ΔN14-CP and WT-CP, we observed a trend of slower spread in case of ΔN14-CP, albeit not statistically significant (Fig. 3B and data S7). ΔN23-CP was not included in the comparison, due to the considerable delay in the systemic spread exhibited by this mutant (Fig. 3A), which made it difficult to measure the viral multiplication area. In conclusion, systemic viral spread is abolished for ΔN26-CP, and decreasingly hampered in case of ΔN23-CP, ΔN19-CP, while ΔN14-CP mutant is able to move similarly as WT-CP, albeit with a short delay.

### Substitution of serine with glycine on position 21 of CP N terminal region abolished PVY cell-to-cell viral movement and virion assembly

To pinpoint amino acids important for cell-to-cell virus movement, we generated point mutations in the part of CP N terminal region that was observed as being important for viral movement (Figs. 1, 2 and 3). We point mutated charged residues Asp14, Glu18, potential phosphorylation site at Ser21 and 3D structure breakers Gly20 and Pro24. All, except Pro24, are highly conserved across sequenced PVY strains. At position 24, 50% of PVY strains carries Pro and the others Ser (Fig S9).

Selected residues were changed to Ala (D14A, E18A, P24A), except for Ser21 where the sequence coding for Ala and Thr substitution was unstable in *E. coli*, thus we mutated it to Gly (S21G) and for Gly20, where we intentionally introduced Pro (G20P) to affect the fold of N terminal region (Fig. 4A).

**Figure 4.**
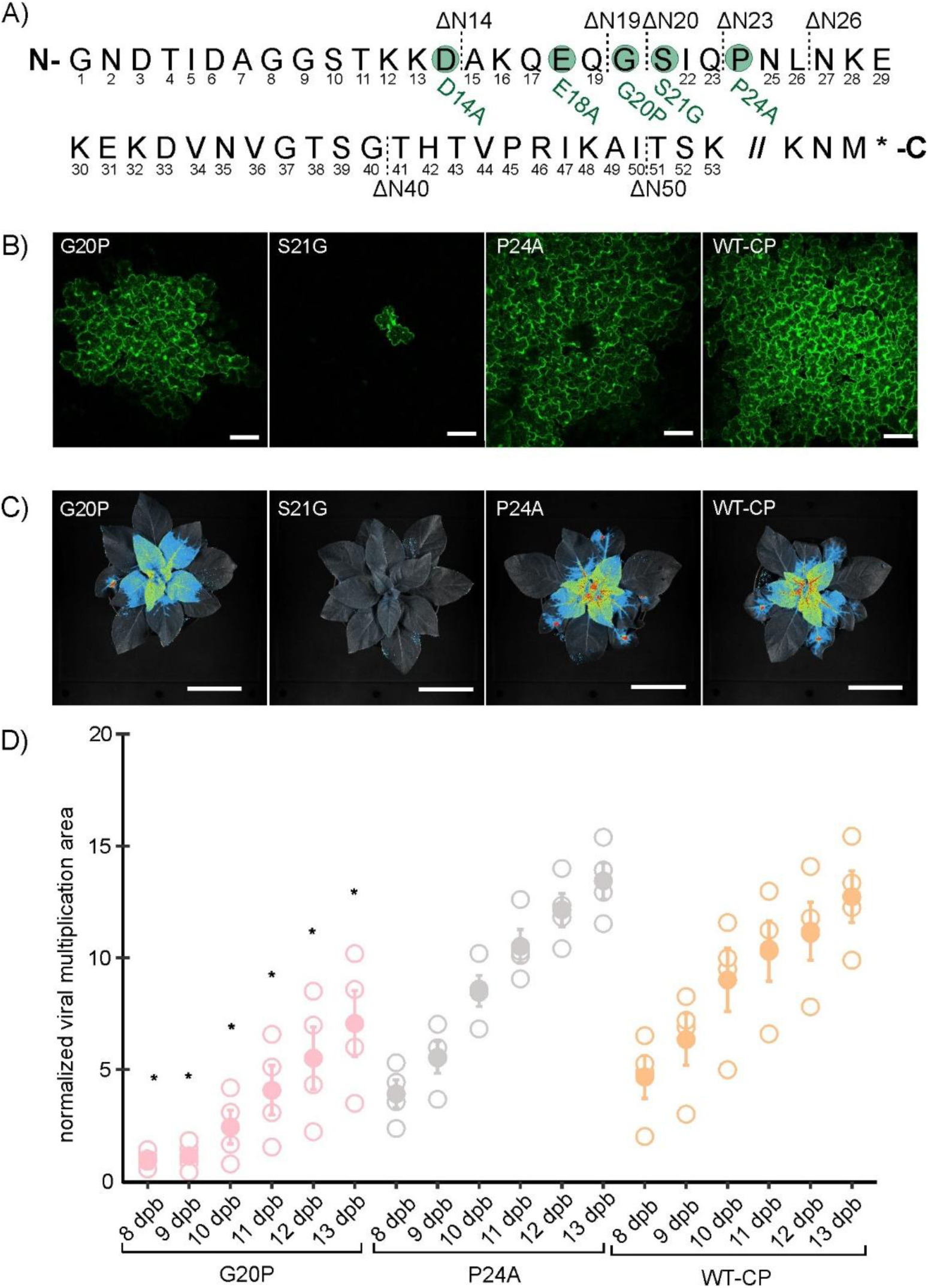
Substitution of serine to glycine on position 21 of CP N terminal region abolished cell-to-cell viral movement. **(A)** Amino acid sequence of GFP tagged infectious clone PVY-N605(123) CP protein with labeled constructed deletion and point mutants. The scheme presents mutants with deletions of 14 (ΔN14-CP), 19 (ΔN19-CP), 20 (ΔN20-CP), 23 (ΔN23-CP), 26 (ΔN26-CP), 40 (ΔN40-CP) or 50 (ΔN50-CP) amino acids on CP N terminal region and mutants with point mutations including substitutions of negatively charged aspartic acid to alanine on position 14 (D14A), negatively charged glutamic acid to alanine on position 18 (E18A), glycine to proline on position 20 (G20P), serine to glycine on position 21 (S21G) and proline to alanine on position 24 (P24A). **(B)** Confocal microscopy images showing viral cell-to-cell spread of mutants G20P, S21G, P24A and WT-CP 5 dpb. Scale is 100 µm. **(C)** Spatio-temporal PVY distribution for constructed point mutants G20P, S21G, P24A and WT-CP in *N. clevelandii* systemic tissue as observed by whole plant imaging system 12 dpb. Scale is 5 cm. **(D)** Quantification of viral multiplication area in the upper systemic leaves of bombarded *N. clevelandii* in 6 selected timepoints (8-13 dpb). Mean (represented with filled dot) and standard error are shown. Individual measurements are shown as empty dots, representing normalized viral multiplication area. Differences between analyzed point mutants were evaluated with Welch’s t test. Statistically significant differences (p<0.05) between G20P mutant and WT-CP are represented with asterisks (*), while there were no statistically observed differences between P24A and WT-CP (see data S8 for all results of statistical testing). Vertical lines present all points except outliers. The results were confirmed in additional experiments (Fig. S5 and data S9).

Viral cell-to-cell spread of mutants D14A, E18A and P24A was comparable to WT-CP (Fig. 4B and S6). On the other hand, G20P mutant showed delayed cell-to-cell spread, while S21G mutant remained limited to just single cells at all observed timepoints (Figs. 4B and S6).

Next, we performed whole plant imaging to study systemic viral spread of mutants (Figs. 4C and S8). In accordance with the results of cell-to-cell movement, there were no statistically significant differences in viral multiplication area in systemic tissue when comparing D14A, E18A or P24A with WT-CP (Figs. 4D, S7, S5 and data S8, S9, S10, S11) at all observed timepoints. On the other hand, the viral multiplication area of G20P in systemic tissue was statistically significantly lower compared to WT-CP at all observed time points (Fig. 4D and data S8). As expected, S21G did not spread systemically (Fig. 4C). Furthermore, viral assembly was confirmed with electron microscopy in all constructed point mutants, with S21G as an exception, where only oligomeric rings were observed (Supp. Fig. S10).

## Discussion

The biological function of C and N terminal regions of PVY CP has already been initiated in our previous study, where we hypothesized that N terminal amino acid residues are necessary for an efficient cell-to-cell movement (17). Here we provide additional insights into the mechanism of PVY spread by identifying the regions of PVY CP N terminal region that are important for efficient viral cell-to-cell movement and systemic infection.

Despite the conserved flexible structure of potyviruses N terminal region of CP, there are notable differences observed regarding its involvement in potyviral cell-to-cell and systemic movement. In the case of zucchini yellow mosaic virus (ZYMV), deletion of the entire CP N terminal region did not affect systemic infection (23). Similarly, a deletion of 6-50 amino acids in the CP N terminal domain did not compromise TuMV cell to cell and systemic movement (19). On the other hand, in tobacco etch potyvirus (TEV), CP N terminal region deletions of 5-29 amino acid residues delayed cell-to-cell movement and prevented systemic spread (22). We observed inability to move cell-to-cell for PVY when 40 or more amino acid residues were deleted from CP N terminal region (Fig. 1). However, shorter deletions in the range of 19 to 23 did not prevent systemic spread but resulted in delayed systemic and cell-to-cell PVY movement (Figs. 2, B and C, 3B and data S4, S3 and S7).The observed differences among potyviral species may stem from the high phylogenetic divergence inherent to the genus, which at the CP level is, among other variations, observed as different lengths and nucleotide variations in the N terminal region across different potyvirus species (29, 23, 30, 31, 32). It is also well known that the potyvirus movement is facilitated by various protein components, both from the virus and the host plant (33, 34, 35, 10, 11). In addition potyviruses exhibit a divergent host range, majority of them being naturally limited to narrow host range, such as TEV and ZYMV (36, 37), while TuMV stands out with its remarkably broad natural host range (38). These features could also be contributing to the differences in the influence that CP N terminal domain deletions have on the movement of different potyvirus species, because different components involved in potyviral movement, viral and host, could be differently compensating the effect of CP N terminal deletions.

The analysis of viral multiplication area revealed important insights into the cell-to-cell movement dynamics of deletion mutants, as increasing number of deleted amino acid residues at CP N terminal region resulted in slower cell-to-cell viral spread (Fig. 2, B and C). Furthermore, slower cell-to-cell virus spread also impacted infection of systemic tissue (Fig. 3A). The connection between cell-to-cell movement and systemic spread was expected, as the virus reaches the veins, enters the veins and is transported in systemic tissue with a delay due to decelerated cell-to-cell movement (39).

Results of several studies showed the role of charged amino acids in the CP core, C and N terminal region of different potyviruses in cell-to-cell and systemic virus spread (20, 26, 40, 41). In contrast, substitutions of charged amino acids in the CP N terminal region of PVY in our study (D14A and E18A mutants) did not affect cell-to-cell spread (Fig. 4B). It was also reported (42) that aromatic residues, located on the core CP region contribute to the formation of π-stackings that are importantly involved in TVBMV and other potyvirus movement, since they maintain CP accumulation by stabilizing α-helixes and β-sheets (43, 44). Our region of interest does not possess aromatic amino acid residues, therefore we did not test this hypothesis.

However, we observed that replication of S21G mutant was limited to single cells (Fig. 4B). We further checked and the intensity of replication of this mutant within this single cell was not affected (Table S2). Interestingly, electron microscopy imaging showed impaired viral assembly, since only oligomeric rings were observed (Supp Fig. S10). Correct viral particle assembly is however not prerequisite for cell to cell movement as ΔN40 and ΔN50 mutants did assemble but still could not move between cells (Figs. 1 and S10). We hypothesize that this might be due to loss of potential phosphorylation site or disruption of N-terminus structure, due to introduction of glycine. Previous studies suggested that posttranslational modifications such as phosphorylations have an important role in viral cell-to-cell movement and virion assembly (45, 46). In potato virus A (PVA), phosphorylation of Thr243 residue on CP C terminal region is crucial for PVA replication in planta (21). Note that to confirm the hypothesis about role of phosphorylation of Ser21, we aimed to replace Ser21 also with alanine and threonine, but the design of such point mutants was not achievable, due to their instability in *E. coli*. It has been previously suggested that PVY cDNA is unstable in *E. coli*. Stability of full length PVY plasmids was achieved by inserting introns into putatively toxic genes but still for some mutants recombinations can occur (47).

Furthermore, we also showed the important role of the glycine on position 20 of CP N terminal region in PVY cell-to-cell as well as systemic spread, since G20P mutant spreads slower than WT-CP (Fig. 4 B, C and D). We hypothesize that exchanging Gly with Pro in G20P mutant increased rigidity in the N terminal region. Consequently, the interaction of the CP with other host and viral proteins in the cell may be impaired, resulting in the observed delay. Moreover, alignment of first 50 amino acid residues from PVY CP N terminal region across all PVY isolates, revealed a high degree of conservation (at least 92,9%) at the mutated residues 20 (G20P) and 21 (S21G) across all PVY isolates (Fig. S9), further supporting biological significance of these amino acids in PVY.

The results identifying crucial domains and amino acids of CP N terminal region in viral cell-to-cell movement provided in our study are important basis to elucidate these complex processes, particularly in most economically important viruses such as PVY.

## Materials and Methods

### Plant material

*N. clevelandii* plants were used to follow virus spread. Plants were grown from seeds and kept in growth chambers under controlled conditions as described previously (48).

### Construction of PVY-N605(123)-GFP CP N-terminal deletion and point mutants

As a template to construct mutants with deletions on the CP N terminal region, we used a GFP-tagged infectious clone PVY-N605(123) (28). The GFP coding sequence was inserted between NIb and CP coding sequences, flanked by protease target sequences, so that GFP would not bother viral movement, since it is excised from the polyprotein as other viral proteins (28). Megaprimers were designed in accordance with instructions prepared by (49). Detailed information about mutagenic PCR reactions conditions and mixtures can be found in Text S1. After adding 4 µL of DpnI enzyme, the mutagenesis mixtures were transformed into *E. coli* XL 10 Ultracompetent Cells in accordance with manufacturer’s instructions (Agilent Technologies). Transformation mixtures were plated on LB agar with ampicillin selection and incubated overnight at 37⁰C. Transformants were screened with colony PCR using PVY GFP_F and PVY uni_R primers (data S12) and KAPA2G Robust HotStart Kit (Agilent Technologies). Prior bombardment, polyprotein and its promoter in PVY-N605(123) lacking 50 and 14 amino acids on CP N-terminus were Sanger sequenced (data S12). Additionally, restriction analysis was performed to confirm the correctness of the PVY coding region.

### Biolistic bombardment

Constructed PVY mutants and wild type plasmids were amplified in *E. coli*. Plasmids were then isolated and coated onto gold microcarriers that were used for *N. clevelandii* bombardment using a Helios® gene gun (Bio-Rad), as described by (49).

### Plant material sampling and RNA isolation

Plant leaf samples were collected from bombarded *N. clevelandii* plants by excising 0,3 g plant tissue near the bombardment site. Sampled plant material from bombarded *N. clevelandii* leaves was homogenized in 700 µl of RNeasy kit RLT lysis buffer with FastPrep (MP Biomedicals) 1 min 6,5 m/s. Total RNA was isolated using RNeasy Plant Mini Kit (Qiagen). Residual DNA was digested with deoxyribonuclease I (DNase I, Qiagen) in solution, using 1.36U DNase/µg RNA. Quality control of isolated RNA was performed as described (48).

### RT-qPCR

Relative PVY RNA concentration was determined by single-step quantitative RT-PCR (RT-qPCR; AgPath-ID One-Step RT-PCR, Thermo FisherScientific) for the ΔN50-CP mutant, while two step RT-qPCR was performed for other mutants. RNA was reverse transcribed with High-Capacity cDNA Reverse Transcription Kit (Thermo Fisher Scientific) and analyzed by qPCR using FastStart Universal Probe Master (Roche).

A qPCR assay targeting the PVY CP encoding region was used as a target gene, while the gene encoding for cytochrome oxidase (COX) was used as reference gene. To avoid false positive results plants bombarded with plasmid encoding for blue fluorescent protein, were sampled and analyzed with our qPCR assay targeting PVY CP encoding region. RT-qPCR amplification program and reaction composition mixtures were the same as previously described (28). Primers and Probes details are listed in data S12.

After conducting RT-qPCR reactions, the QuantGenius software (50) was used for relative quantification of PVY RNA using the standard curve approach. Data containing information about the relative RNA quantity, was averaged for each group and then normalized to the lowest group average number (data S1).

### RT-PCR

One-step reverse transcription PCR (QIAGEN OneStep RT-PCR Kit) was performed to confirm that the introduced mutations were maintained in the viral progeny of all constructed mutants. DNAse-treated RNA, isolated from leaves according to the above protocol, was used as a template. Mutated region was amplified with PVY-GFP_F and uniPVY_R primers (data S12). The purified PCR products were sequenced (Sanger sequencing). For samples with a small amount of mutated virus (ΔN50-CP, ΔN40-CP, S21G), an additional PCR (repliQa HiFi ToughMix; Quantabio) on purified products was performed to amplify the mutated region in sufficient quantity.

### Confocal microscopy

Cell-to-cell viral spread on upper side of bombarded *N. clevelandii* leaf discs, sampled with cork borer, was observed under Leica TCS LSI confocal macroscope with Plan APO x5 and x20 objectives (Leica Microsystems, Wetzlar, Germany) at different time points (5-14 dpb), and Stellaris 5 confocal microscope with HC PL FLUOTAR x10 objective (Leica Microsystems, Wetzlar, Germany) for studying point mutations at 5 dpb. The GFP emission was recorded after excitation with 488 nm laser in the window between 505 and 530 nm and used as a direct measure of the presence of metabolically active viruses in the observed tissue. Sampled discs were scanned unidirectionally with scan speed of 400 Hz and frame average 1 or 2. For following the spread of ΔN50-CP at 12 dpb, 5x and 20x objective were used (Fig. 1B), for ΔN40-CP and WT-CP at 5 and 10 dpb, 5x objective was used (Fig. 1B), while for following the spread of ΔN26-CP at 5-14 dpb and ΔN23-CP, ΔN20-CP, ΔN19-CP, ΔN14-CP mutants and WT-CP at 5 dpb, 20x objective was used (Fig. 1C and D). For each mutant we observed at least 3 plants.Images were processed using LEICA LAS X software (Leica microsystems) to obtain maximum projections from z-stacks. Raw confocal microscopy images were deposited on Zenodo (doi: 10.5281/zenodo.17177382).

### Transmission electron microscopy sample preparation

Viral assembly was assessed with transmission electron microscopy (TEM) using negative staining method. Tissue extracts were prepared from leaf discs, in which viral presence was previously confirmed with confocal microscope, and then macerated in phosphate buffer (0.1 M, pH 6.8). 20 µL of extracts were applied to a grid for 5 minutes, blotted, washed, and stained with 1% (w/v) water solution of uranyl acetate. Grids were observed by TEM TALOS L120 (Thermo Fisher Scientific) operating at 100 kV or 120 kV. Micrographs were recorded using Ceta 16 M camera and software Velox (Thermo Fisher Scientific). Raw electron microscopy images were deposited at Zenodo (doi: 10.5281/zenodo.17177382).

### Virus inoculation

To more precisely examine cell-to-cell viral movement by whole plant imaging, we inoculated 3-4 weeks old *N. clevelandii* plants with inoculum prepared from systemic leaves of ΔN14-CP, ΔN19-CP, ΔN23-CP and WT-CP-inoculated plants in which GFP signal was confirmed using whole plant imaging system. Systemic leaves were ground in phosphate buffer (supplemented with PVP 10000) in a plant material: buffer ratio of 1:4. Three bottom leaves of *N. clevelandii* plants were dusted with carborundum powder and rubbed with the inoculum (1-3 drops of inoculum per leaf) which was removed after 10 min by rinsing with tap water.

### Whole plant imaging

Cell-to-cell and systemic viral spread throughout the plant was monitored using the Whole plant imaging system Newton 7.0 BIO (Vilber) in bombarded and inoculated *N. clevelandii* plants. GFP emission of GFP-tagged PVY clone was followed in 480 nm excitation channel and emission filter F-550. Images were taken by EvolutionCapt edge software. For viral multiplication area analysis in inoculated leaves, all three ΔN14-CP-, ΔN19-CP-, ΔN23-CP- and WT-CP-inoculated leaves of *N. clevelandii* were imaged using exposure time 1 min 46 sec 700 msec, 10×10 field of view (FOV) and 1331-1337 focus in 4 time points: 3 dpi a.m. and p.m. and 4 dpi a.m. and p.m. Measurements for a.m. timepoint were carried out 9 a.m, while for p.m timepoint at 1 p.m. Four plants inoculated with each mutant or WT-CP were imaged. For studying systemic viral spread, at least 3 *N. clevelandii* plants bombarded with ΔN26-CP, ΔN23-CP, ΔN14-CP, ΔN19-CP- and WT-CP were imaged, using 50 s exposure time, 20×20 FOV and focus in the range between 1871 and 1905 in different time points (7-12 dpb). Note that *N. clevelandii* plants bombarded with ΔN26-CP and ΔN23-CP were not taken into analysis, due to their inability or big delay in systemic movement. For studying systemic viral spread of point mutations, at least three plants bombarded with constructed point mutants were imaged at different time points (6-13 dpb) with exposure time 50 s and other settings as for studying systemic viral spread of deletion mutants (see above). In case of experiment with D14A and WT-CP at 13 dpb, exposure time was only 5 s to avoid saturation due to a high signal. Raw images were deposited at Zenodo (doi: 10.5281/zenodo.17177382).

### Image analysis and signal quantification following whole plant imaging

Viral multiplication area analysis on inoculated leaves and on upper systemic leaves were performed using Kuant software (Vilber, France). In case of viral multiplication area, surfaces of three viral multiplication areas per leaf were measured, while in case of viral multiplication area in systemic leaves, total count of the virus affected area was measured. Total count is determined as a sum of gray values within region of interest (systemic leaves, ROI) and is proportional to the intensity and spread of the signal. All measured data from Kuant software were exported to Excel, where total count in ROIs were averaged for each timepoint and subsequently normalized on the lowest group average number. Normalized viral multiplication areas were statistically evaluated using Welch’s t-test and were used for graphic representation.

## Acknowledgments

We thank Živa Lengar for technical support and laboratory assistance, Dr. Maja Zagorščak and Dr. Angelika Vižintin for help with statistical analysis.

## Author contributions

AC, KG and TL designed research; TMP, KS, AC, AV, MTŽ, KB, VL performed the research; AV, TL, TMP, KS, MTŽ, VL, KG, AC, analyzed the data; AV, TL, AC, KG, TMP, KS, MTŽ, IGA, MP, KB, VL contributed to the writing or revision of the article.

## Competing interests

The authors declare no competing interests.

## Data availability

Raw confocal microscopy and whole plant imaging system pictures were deposited at Zenodo and are openly available at doi: 10.5281/zenodo.17177382).

## Supporting Information for

### Supporting Information Text S1. Construction of PVY-N605(123)-GFP CP N-terminal mutants

Mutants were prepared with mutagenic PCR using QuikChange II XL Site-Directed Mutagenesis Kit (Agilent Technologies). As a template previously constructed GFP infectious clone PVY-N605(123) was used (28). The megaprimers were synthesized using the N-terminal region sequence of GFP tagged PVY-N605(123) plasmid, according to defined guidelines (49). All megaprimers used in the study are listed in data S12. Mutagenic touchdown PCR reaction program with the following reaction mixture in the final volume of 25 µL were the same for all generated mutants according to previously published protocol for generation of PVY deletion mutants (49), with minor modifications listed below.

| Component | Final concentration | Volume [µL] |
| --- | --- | --- |
| 10x reaction buffer | 1x | 2,5 |
| dNTP mix | 200 µM | 0,5 |
| forward megaprimer | 0,5 µM | 1,25 |
| reverse megaprimer | 0,5 µM | 1,25 |
| QuikSolution | / | 1,5 |
| PVY-N605(123) | 200 ng | 1,2 |
| PfuUltra HF DNA polymerase | / | 1 |

| Temperature | Time | Step |
| --- | --- | --- |
| 92°C | 2 min | hold |
| 92°C | 50 s |  |
| 65°C to 55°C | 50 s | 10cycles |
| 68°C | 30 min* |  |
| 92°C | 50 s |  |
| 55°C | 50 s | 8 cycles |
| 68°C | 30 min* |  |
| 68°C | 60 min | hold |

After amplification, 4 µL of DpnI enzyme (Agilent Technologies) was added to the mutagenesis reaction mixture, following 2h on 37⁰C incubation. Following DpnI digestion, 2 µL of mutagenesis mixture was used for transformation into *E. coli* XL-10 Gold Ultracompetent Cells (Agilent Technologies). We used 45 µL cell aliquot supplemented with 2 µL of β-mercaptoethanol for the standard heat-shock transformation protocol in accordance with the manufacturer’s instructions (Agilent Technologies). Transformation mixtures were plated on LB agar containing ampicillin and incubated overnight at 37⁰C. Transformants were analyzed with colony PCR using primers PVY GFP_F and PVY uni_R with KAPA2G Robust HotStart Kit (Agilent Technologies) with the following 10 µL reaction mixture and cycling conditions stated below.

| Component | Final concentration | Volume [µL] |
| --- | --- | --- |
| 5x Buffer B | 1x | 2 |
| 10 mM dNTP | 200 µM | 0,2 |
| 10 µM PVY GFP_F | 300 nM | 0,3 |
| 10 µM PVY uni_R | 300 nM | 0,3 |
| 5 U/µL KAPA2G polymerase | 0,3 U | 0,06 |
| Colony suspension | / | 1 |
| H2O |  | 6,14 |

| Temperature | Time | Step |
| --- | --- | --- |
| 95 °C | 10min | Hold |
| 95 °C | 30 s |  |
| 55 °C | 15 s | 30 cycles |
| 72 °C | 1 min |  |
| 72 °C | 5 min | Hold |

Sanger sequencing, using the same primers as for the colony PCR, of positive colonies was performed to confirm correct sequence of the PVY coding part.

Designed mutants PVY-N605(123)-GFP with desired mutations were amplified in One Shot® TOP10 *E. coli* and 50 µg of constructed plasmid mutants were isolated from overnight cultures using GenElute Plasmid MiniPrep Kit (Sigma-Aldrich). Isolated plasmids were subsequently used to coat 6.25 mg of gold microcarriers (0,6 µm) to prepare gene gun bullets according to the manufacturers protocol and were used for *Nicotiana clevelandii* bombardment using a Helios®gene gun (Bio-Rad) at 200 psi (49).

## Supporting figures and tables

**Fig. S1.**
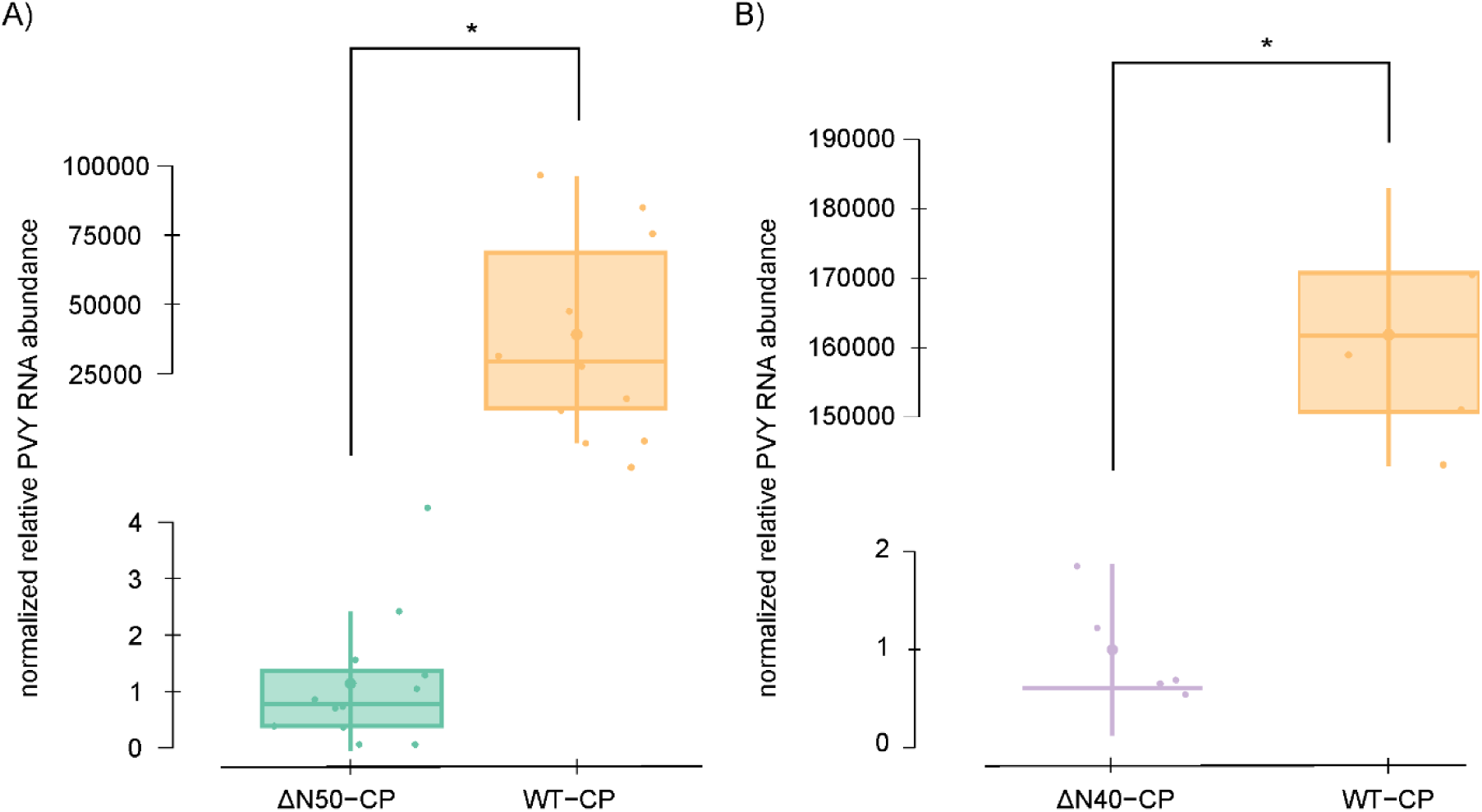
ΔN50-CP ΔN40-CP viral replication. Normalized relative PVY RNA abundance in bombarded *N. clevelandii* leaves for constructed PVY mutants lacking 50 (ΔN50-CP) (A) and 40 (ΔN40-CP) (B) amino acids at CP N-terminus. Results were obtained 14 days post bombardment (dpb). Non-mutated infectious clone (WT-CP) was used as a control. Data normalization was performed as described in data S1. Results are presented as boxplots with normalized relative PVY RNA abundance for each sample shown as dots. Differences between ΔN50-CP and WT-CP and between ΔN50-CP and WT-CP were statistically evaluated using Welch’s t test. Statistically significant differences (p<0,05) are marked with an asterisk (*). Vertical lines present all points except outliers.

**Fig. S2.**
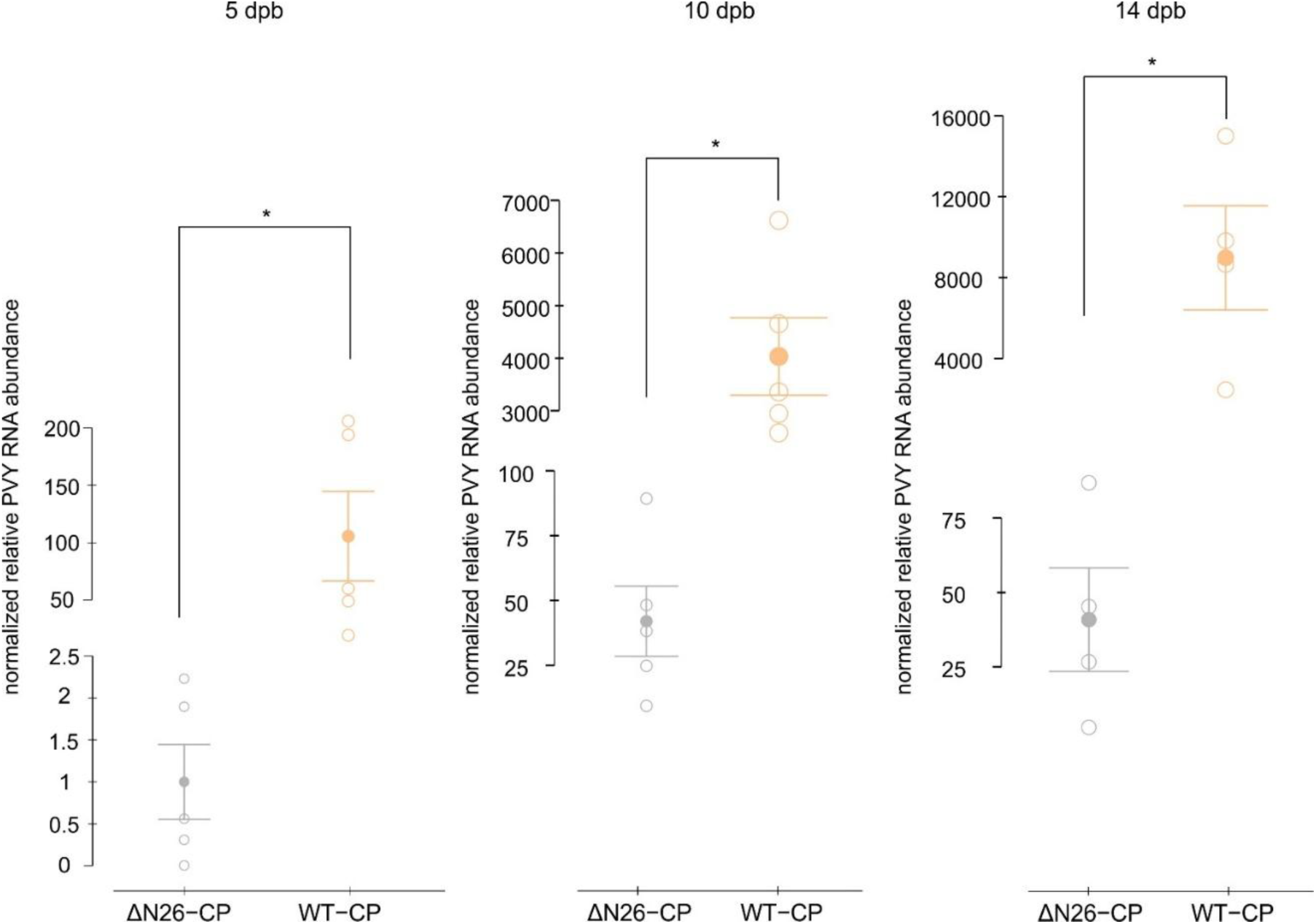
Viral replication ΔN26-CP. Normalized relative PVY RNA abundance in bombarded *N. clevelandii* leaves for constructed PVY mutant lacking 26 amino acids residues, in three timepoints including 5, 10 and 14 dpb (from left to right). Non-mutated infectious clone (WT-CP) was used as a control. Data normalization was performed as described in data S1. Results are presented as mean (represented with filled dot) and standard error. Individual measurements are shown as empty dots, representing normalized relative PVY RNA abundance. Differences between constructed deletion mutants and WT-CP were statistically evaluated using Welch’s t test. Statistically significant differences (p<0,05) are marked with an asterisk (*). Note that the scales are different between time points.

**Fig. S3.**
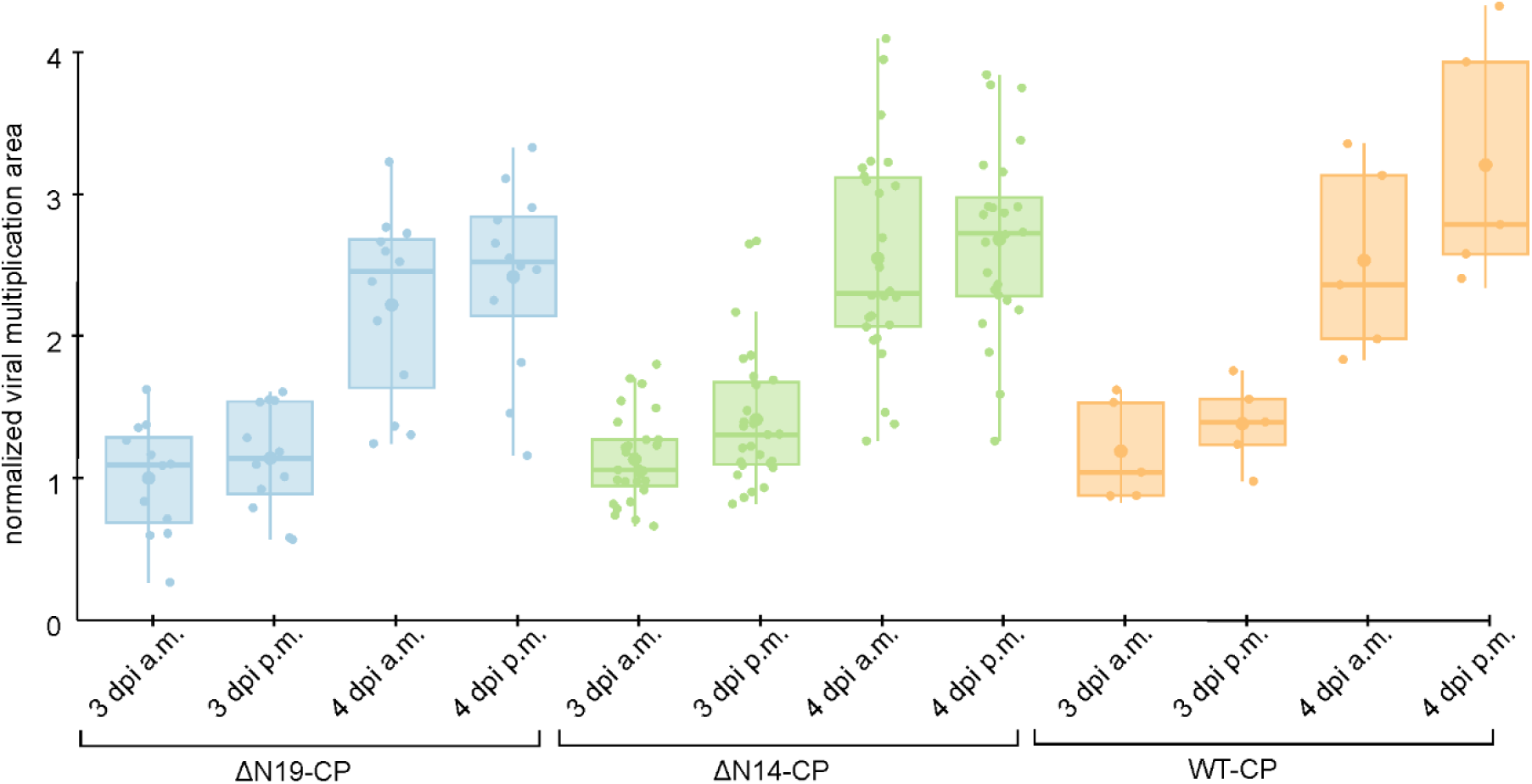
N19-CP_ΔN14-CP_WT-CP cell-to-cell spread dynamics. Viral cell-to-cell spread dynamics was quantified with normalized viral multiplication analysis as described in Materials and methods. Results are presented as boxplots for tested mutants ΔN19-CP, ΔN14-CP and WT-CP in 4 tested timepoints including 3 dpi a.m., 3 dpi p.m., 4 dpi a.m. and 4 dpi p.m., where dots are representing normalized viral multiplication area as described in data S5. Differences were statistically evaluated using Welch’s t test. Vertical lines present all points except outliers. The differences were not statistically significant, due to autofluorescence of trichomes which resulted in saturated pixels.

**Fig. S4.**
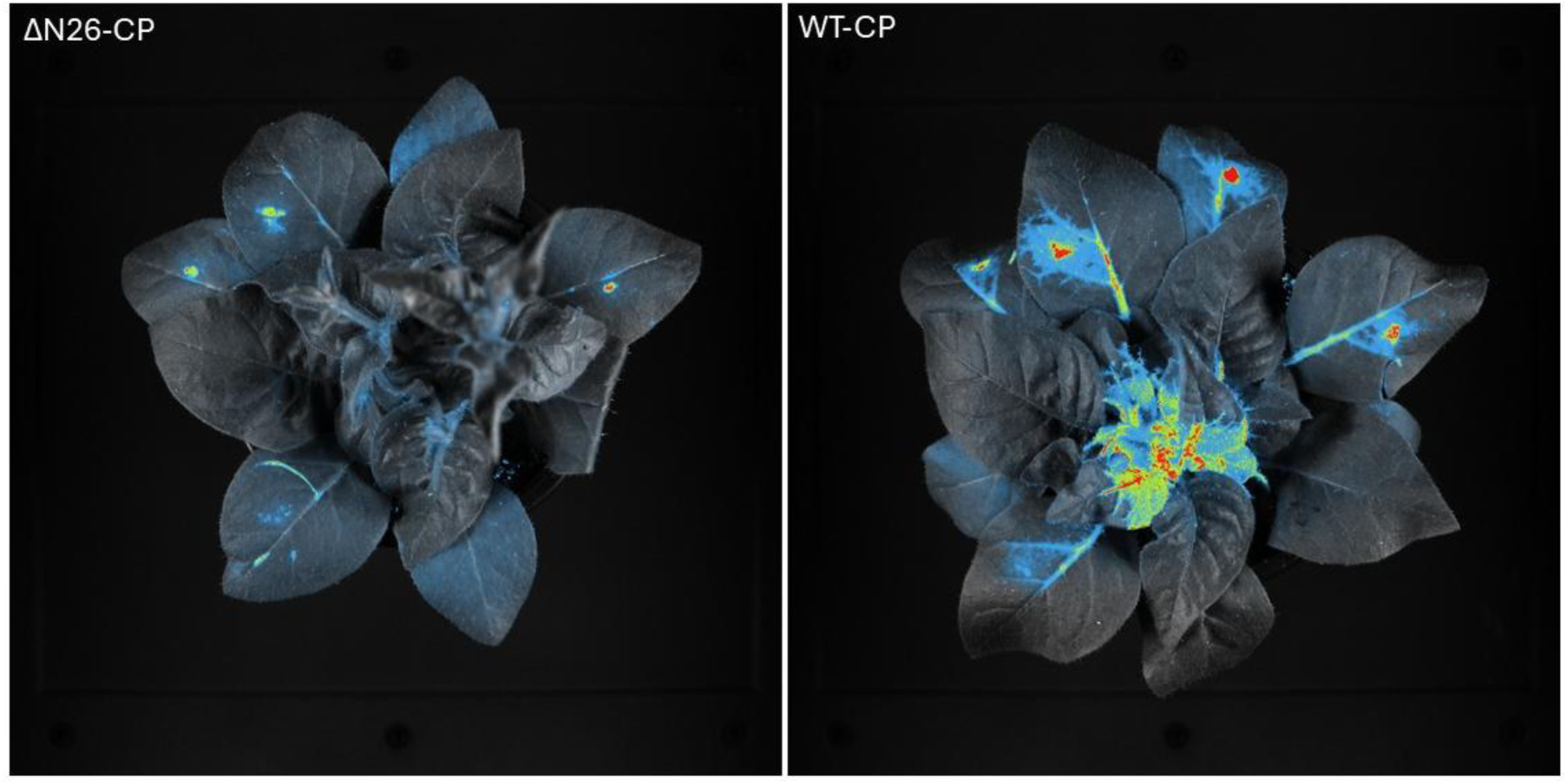
ΔN26-CP inability of systemic spread. Viral systemic spread was abolished in ΔN26-CP mutant (left) in comparison to non mutated WT-CP, where systemic spread occurred (right). Pictures taken 12 dpb.

**Fig. S5.**
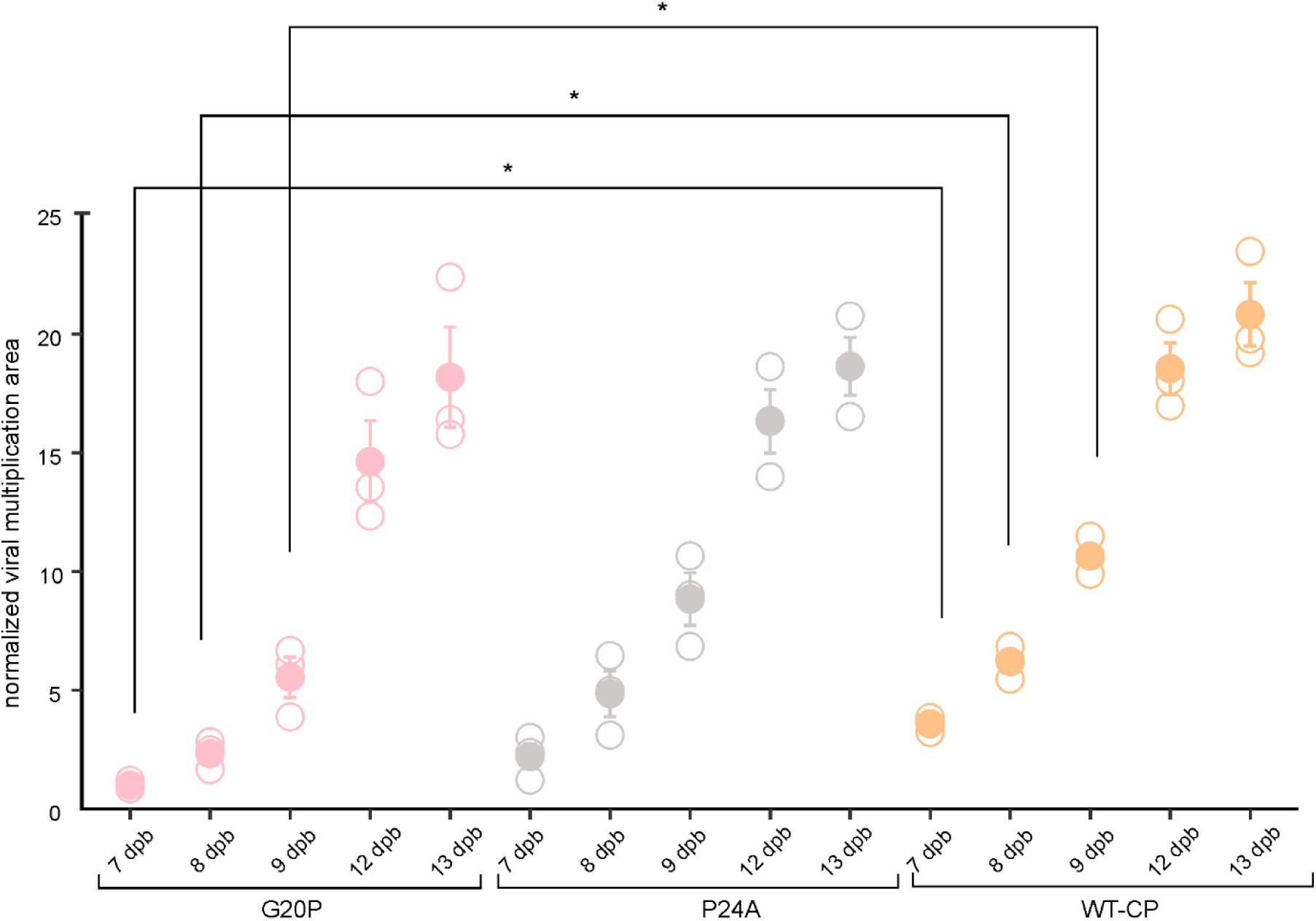
Independent experiment for point mutants G20P, P24A and WT-CP to check viral abundance. Quantification of virus abundance in the upper leaves of bombarded *N. clevelandii* expressed as total count per mutant 7, 8, 9, 12 and 13 dpb with exposure time 50 s (A). Other measurements settings are the same as stated in Materials and methods. Mean (represented with filled dot) and standard error are shown. Individual measurements are shown as empty dots, representing normalized viral multiplication area. Statistically significant difference in normalized viral multiplication area between mutants was evaluated by Welch’s t test. Statistically significant differences (p<0,05) are demarked with an asterisk (*). Raw and normalized data, number of plants and results of statistical analysis are specified in data S9. Note that there was no statistically significant difference between G20P and WT-CP at 12 and 13 dpb due to signal oversaturation due to high exposure time (50 s).

**Fig. S6.**
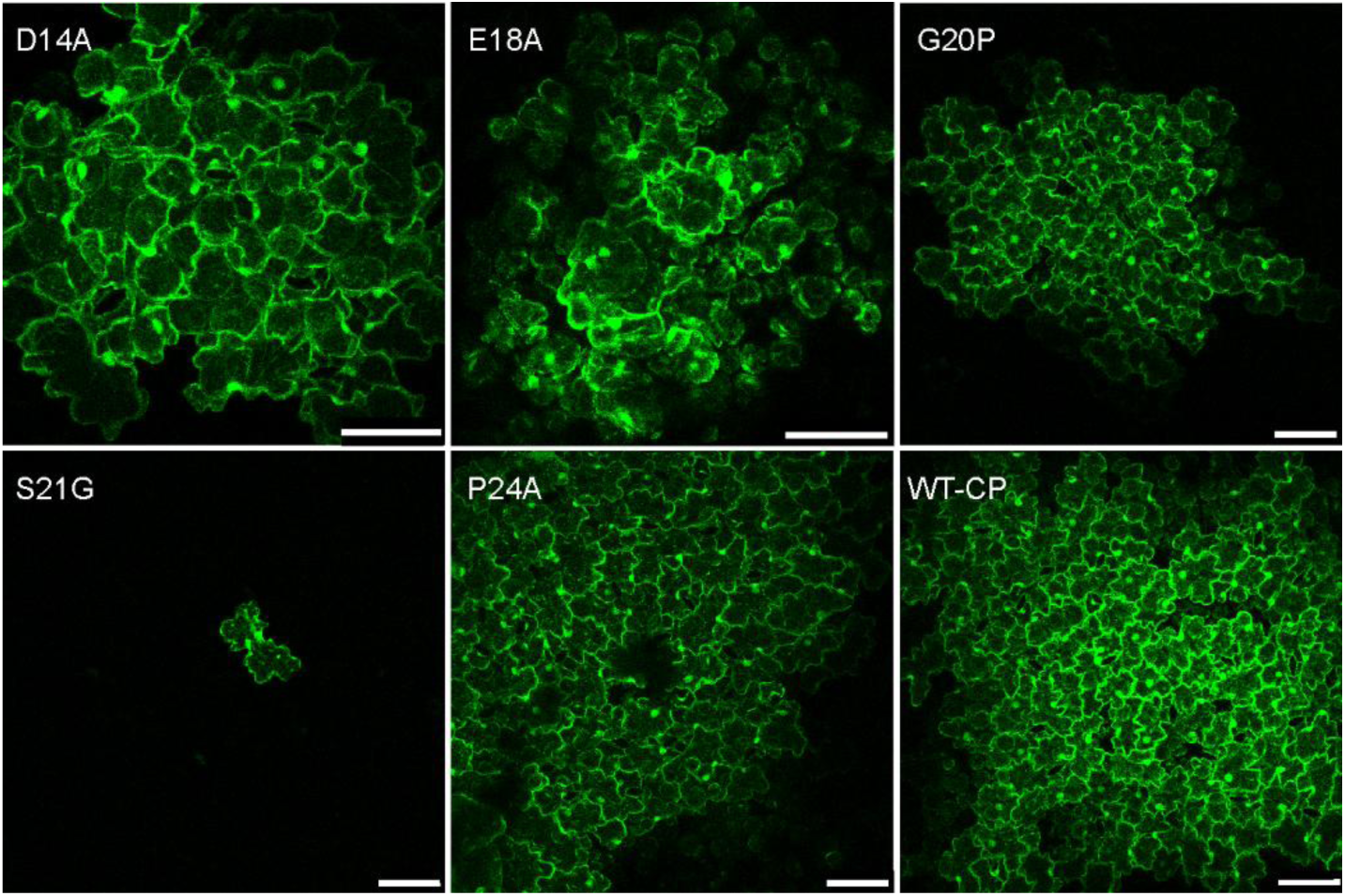
Cell-to-cell viral spread of D14A, E18A, G20P, S21G and P24A point mutants. Confocal microscopy images showing viral cell-to-cell spread of D14A, E18A, G20P, S21G, P24A point mutants and WT-CP 5 dpb. Note that there we have comparisons of D14A and E18A point mutants with the others (G20P, S21G, P24A) already included in the article main text (Fig. 4B). Other confocal microscopy settings are specified in Materials and methods.

**Fig. S7.**
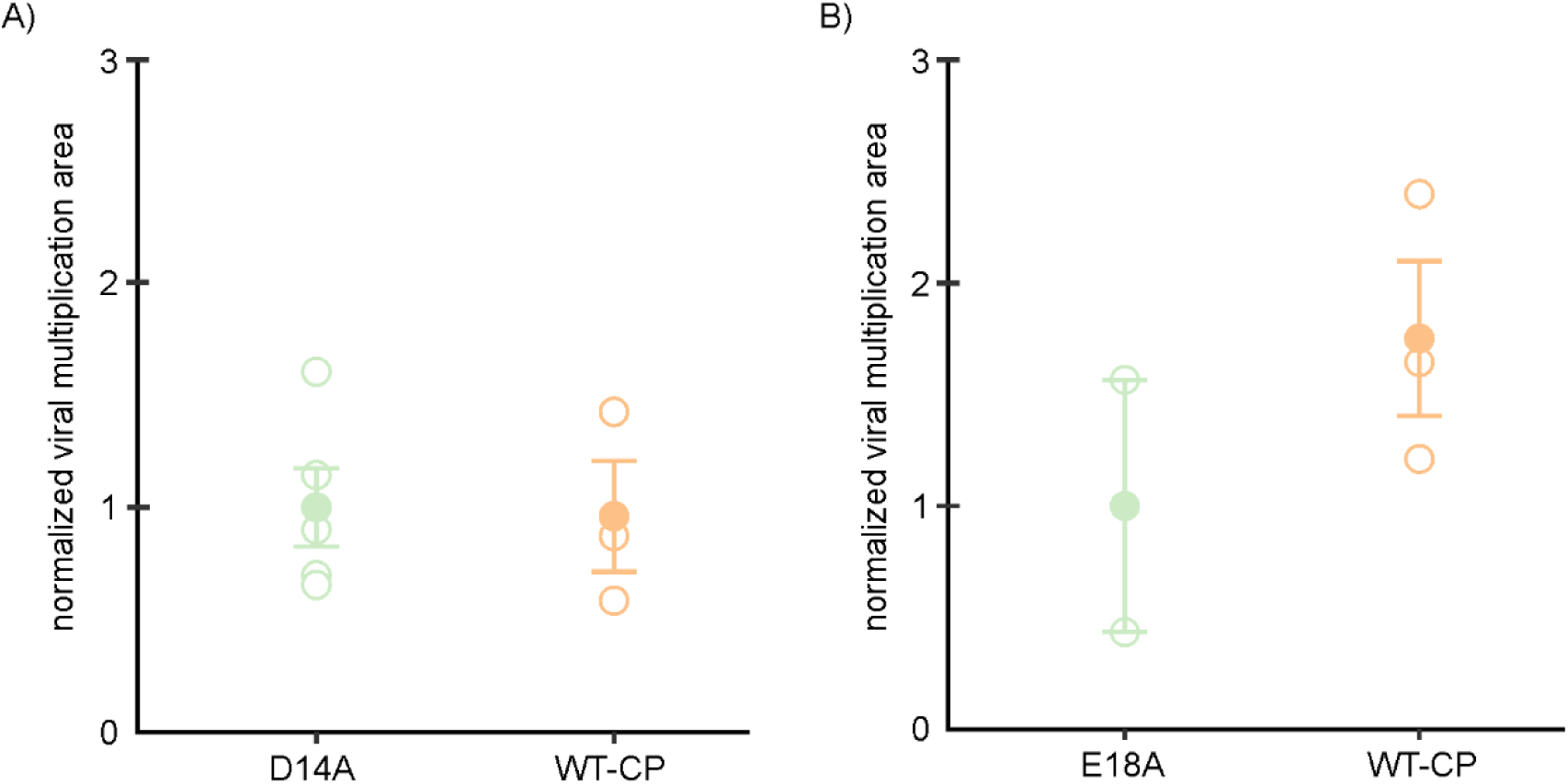
Virus abundance D14A, E18A, WT-CP. Quantification of virus abundance in the upper leaves of bombarded *N. clevelandii* expressed as total count per mutant for D14A mutant 7 dpb with exposure time 50s (A) and (B) for E18A mutant 7 dpb with exposure time 50 (s). Other measurements settings are the same as stated in Materials and methods. Mean (represented with filled dot) and standard error are shown. Individual measurements are shown as empty dots, representing normalized viral multiplication area. There was no statistically significant difference in normalized viral multiplication area between mutants evaluated by Welch’s t test. Raw and normalized data, number of plants and results of statistical analysis are specified in data S10 and S11).

**Fig. S8.**
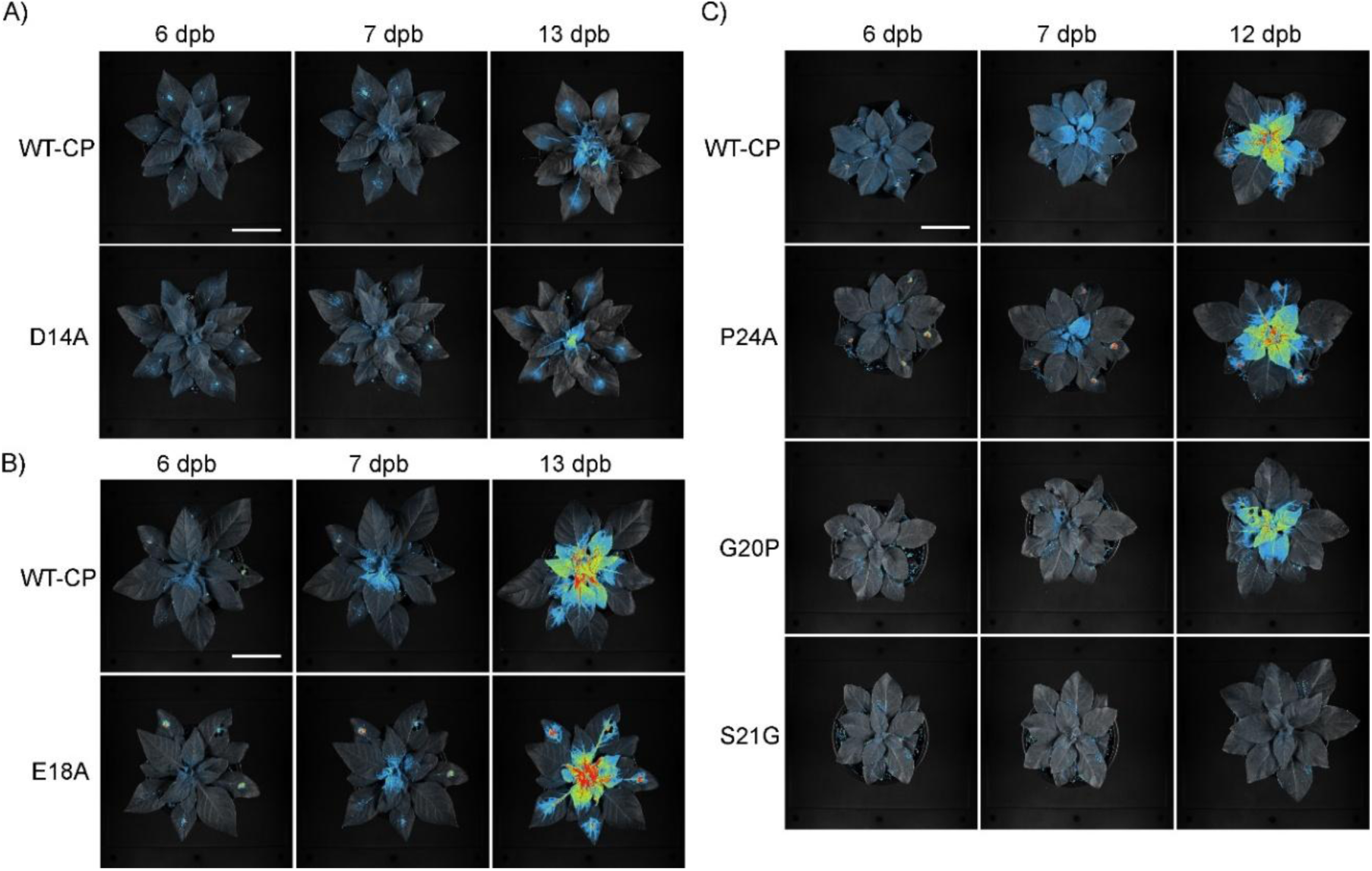
Point mutants systemic spread. Spatio-temporal PVY distribution in *N. clevelandii* systemic tissue using whole plant imaging system. Systemic spread of constructed point mutants was followed 6 dpb, 7 dpb and 13 dpb in case of D14A (A) and E18A (B), while systemic spread of P24A, G20P and S21G. (C) was followed 6 dpb, 7 dpb and 12 dpb. Note that pictures for P24A, S21G and WT-CP 12 dpb are the same as in the main text (Fig. 4C). Imaging settings are specified in Materials and methods. Plants were imaged with exposure time 50 s. In case of D14A and WT-CP at 13 dpb, exposure time was 5 s to avoid oversaturation due to a higher signal.

**Fig. S9.**
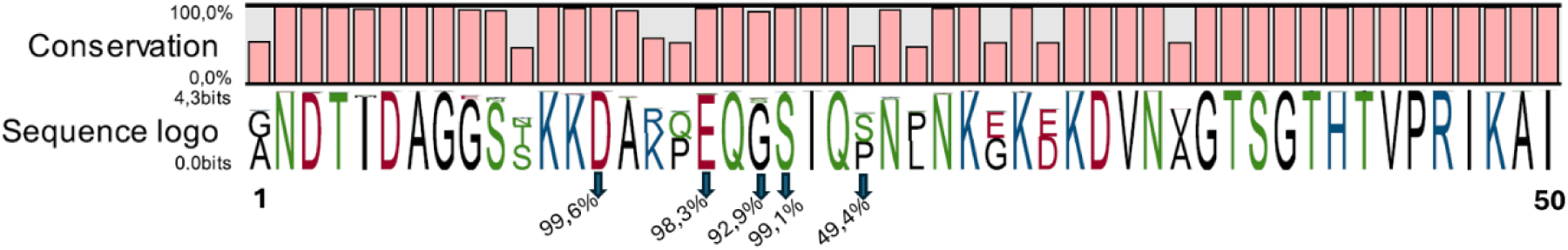
Alignment of first 50 amino acid residues from the PVY N terminal region across all PVY isolates. To assess the conservation of mutated amino acid residues D14A, E18A, G20P, S21G and P24A, we performed multiple sequence alignment of the first 50 amino acid residues of the PVY coat protein, using CLC Genomics Workbench 25 (QIAGEN, Hilden, Germany) and pairwise sequence alignment. Note that for the multiple alignment analysis, only complete sequences containing the first 50 amino acid residues of the PVY coat protein N terminal region were included, as this region corresponds to our engineered deletion and point mutants. Altogether 2112 sequences were obtained from NCBI Virus database. Sequence logo represents the amino acid sequence conservation in the mutated region with arrows showing the conservation percentage of each point mutated amino acid (D14A, E18A, G20P, S21G, P24A) across all obtained sequences.

**Fig. S10.**
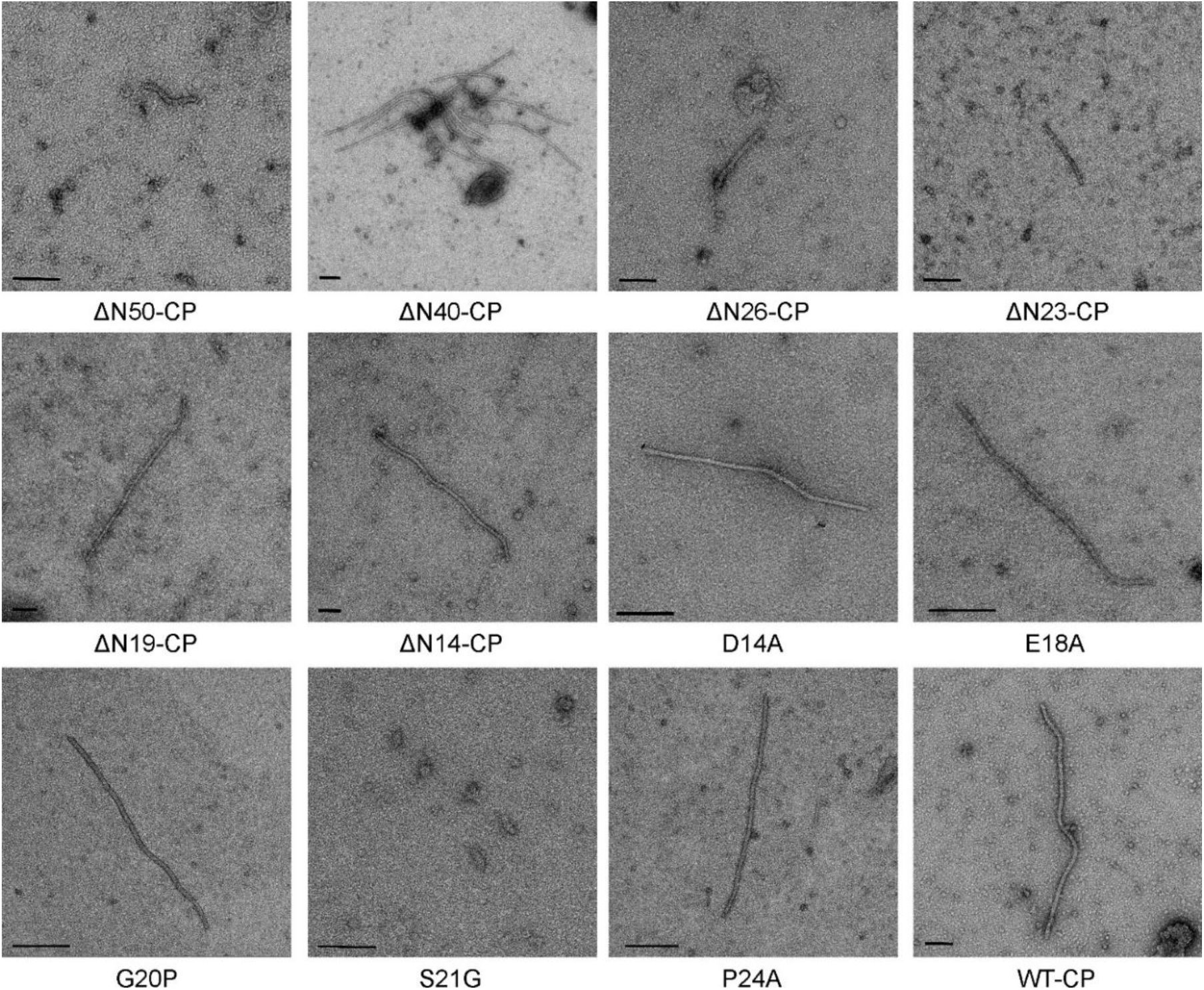
Transmission electorn microscopy (TEM) micrographs of deletion and point mutants. Representative TEM micrographs of deletion mutants and point mutants. Results were obtained with negative staining. Scale bars for deletion mutants and WT-CP are 100 nm and for point mutations 200 nm except in case of S21G (50 nm). Additional images of the mutant viruses were deposited at Zenodo (doi: 10.5281/zenodo.17177382).

---

**Table S1.**
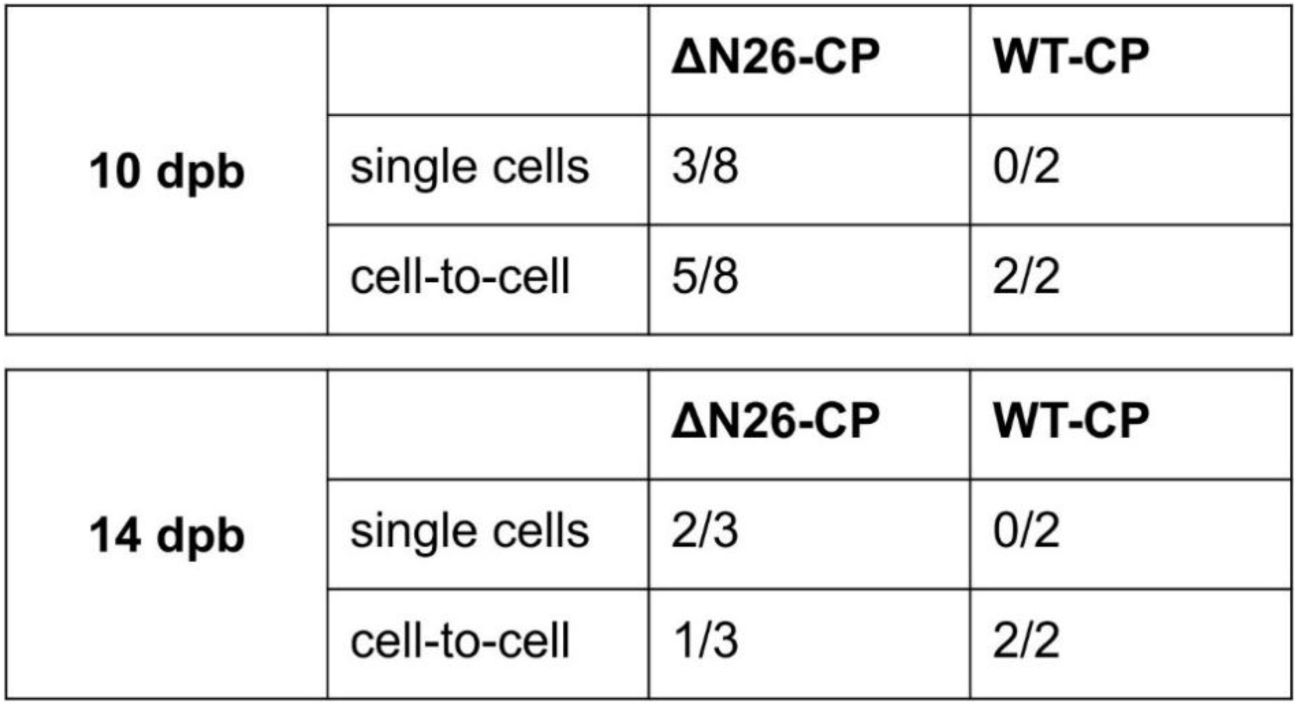
ΔN26-CP viral limitation on single cells or cell-to-cell spread. Number of plants with viral cell-to-cell spread or viral limitation to single cells 10 and 14 dpb. Note that 5 dpb virus was limited to single cells in all observed plants.

**Table S2.** Replication efficiency of ΔN40-CP and S21G mutant is the same as the one of WT-CP PVY. To confirm that detected fluorescent signal in ΔN40-CP and S21G PVY mutants, was the consequence of viral replication and not the continuous expression of viral genes from the original plasmid of PVY driven by the constitutive 35S promoter, ROI (regions of interests) mean intensities of individual cells in confocal microscopy images were assessed. Mean intensities in selected ROIs 5 dpb for ΔN40-CP (A) and S21G (B) compared to WT-CP PVY are shown. Statistical significance of differences was evaluated using Welch’s t test. Note that all images were taken using the same settings (objective, zoom, gain).

| **ΔN40-CP** | **WT-CP** | **Welch`s**<br>**t test** |
| --- | --- | --- |
| 24.4 | 13.8 | 0.6 |
| 14.0 | 17.1 |  |
| 9.7 | 15.5 |  |
| 13.3 | 9.4 |  |
| 8.4 | 15.2 |  |
| 4.8 | 21.6 |  |
| 11.5 | 13.7 |  |
| 8.6 | 15.6 |  |
| 13.1 | 8.7 |  |
| 25.7 | 15.9 |  |
| **13.3** | **14.6** | **average** |

B)
| **S21G** | **WT-CP** | **Welch`s**<br>**t test** |
| --- | --- | --- |
| 6.8 | 7.1 | 0.2 |
| 6.8 | 9.4 |  |
| 6.8 | 12.8 |  |
| **6.8** | **9.8** | **average** |

## Notes

https://zenodo.org/records/17177382

### Competing Interest Statement

The authors have declared no competing interest.

### Summary of Updates

Improved version of the paper with some structural analysis data and in depth discussion.

### Funding

- Slovenian Research Agency: P4-0165, P4-0407, J1-2467

